# Leadership under risk: male vervet monkey’s roles in group progression across high-risk terrain

**DOI:** 10.1101/2025.11.21.689472

**Authors:** Josefien A. Tankink, Erica van de Waal, Redouan Bshary, Carel P. van Schaik

## Abstract

The evolution of cooperative behaviour among unrelated individuals is still incompletely understood. One example is the reported service provisioning by males in primate societies, which is difficult to explain by traditional reciprocity-based models. In this study, we investigated group progressions and sex roles in wild vervet monkeys (*Chlorocebus pygerythrus*) to test two hypotheses for the observed male bias in cooperative and protective behaviours: paternal care versus reputation-based partner choice. Over more than two years, we recorded the order and timing of individual progressions in five neighbouring groups of vervet monkeys across two terrain types differing in risk (rivers and roads). Males, particularly dominant males, were more likely to lead than females or juveniles, and their probability of leading increased in high-risk terrain. When leading, males often crossed alone, thereby assuming higher predation risk, whereas when not leading, males usually crossed within subgroups. Dominant males who had no offspring yet were most likely to lead during risky progressions. Males who did not lead did not preferentially cross with infants based on their paternal status, refuting the paternal care hypothesis. Instead, male leadership in risky contexts appears to represent a cooperative service. While some of these findings are consistent with the reputation-based partner choice model, we found no clear evidence that such services translated into direct mating advantages. Overall, our results highlight the role of dominance and reproductive status in shaping male service provisioning and underscore its potential adaptable function in primate social organisation.

## Introduction

In primate societies, males frequently assume more active roles than females in predator avoidance by engaging in costly behaviours such as increased vigilance, mobbing, and even attacking predators (Baldellou et al., 1992; Stephan & Zuberbühler, 2016; Stojan-Dolar & Heymann, 2010; van Schaik & van Noordwijk, 1989). These behaviours can be considered helping acts, since they are costly for the acting individual but beneficial for the rest of the group (Baldellou et al., 1992; Roberts, 1998; van Schaik & van Noordwijk, 1989; van Schaik et al., 2022). This sex-bias in cooperative behaviour, also called *male services*, has been found to be widespread across primate societies and seems to be independent of the social system (van Schaik et al., 2022). Importantly, there is little evidence that such services are reciprocated with short time delays by other group members. For example, in vervet monkeys, male contributing to fighting neighbouring groups may receive some grooming by females (Arseneau et al. 2018), but that can hardly compensate for the risk of injury. Thus, male services do not fit assumptions about reciprocal exchanges of services as modelled in standard stylised games such as the iterated prisoner’s dilemma (Axelrod & Hamilton 1981). Instead, male services provide opportunities to investigate how cooperative behaviour is affected by key biological variables, such as rank and life history state.

The roles of males in group movement provides a suitable context to study male services in more detail. According to the “selfish herd” model (Hamilton, 1971), predation risk shapes socio-spatial positioning, with central positions being safer but more competitive. Directional movement adds another gradient: individuals at the front face higher risks of encountering predators first (Bumann et al., 1997). To adapt to these gradients of risk, group-living animals may adjust their behaviour, leading to potential differences in roles and positioning of group members, based on individual characteristics. When predation risk is relatively low, females are generally known to lead group progressions in primates (Barelli et al., 2008; Fichtel et al., 2011; van Schaik et al., 2022). Yet, when predation risk increases, males have been shown to place themselves in more vulnerable positions, also called inter-positioning, such as at the front or the back of the group (Hockings, 2011; Tellier et al., 2025). This inter-positioning, which increases individual predation risk, can be considered a service to other group members, i.e. a cooperative behaviour (Hockings, 2011; Tellier et al., 2025; van Schaik et al., 2022). Although much research has gone into spatial positioning and the corresponding risk (for an overview, see Suire et al., 2023), the specific roles of different group members, particularly in dynamic situations and across varying risk environments, remain a topic of ongoing investigation (Suire et al., 2023).

One hypothesis that could explain the sex-bias in this inter-positioning behaviour posits that male service provisioning represents a form of paternal care (van Schaik et al., 2022). In many primate societies, where males and females are continuously associated in multimale/multifemale groups (Cheney et al., 1986; van Schaik & Paul, 1996), males may enhance their inclusive fitness by improving the survival prospects of their offspring – a prediction derived from Hamilton’s rule (Hamilton, 1964). However, this cannot explain cases where newly immigrated males – who lack offspring – also provide services (Baldellou et al., 1992; Matsumoto-Oda et al., 2018; Stephan & Zuberbühler, 2016).

An alternative explanation for the prevalence of male services is reputation-based partner choice (Roberts, 1998; Roberts et al., 2021; van Schaik et al., 2022). Here, individuals perform helping acts to signal their quality and secure future mating opportunities – a process conceptualized as a handicap, where only individuals of high quality can afford to incur the cost of cooperation (Roberts, 1998; Zahavi, 1975). When there is reputation-based partner choice, having a cooperative or helping reputation should increase the male’s benefits in other contexts, especially mating opportunities. However, similar behaviour can be expected when males perform services as an investment into female survival, which should ultimately benefit their mating opportunities (Bshary et al., 2022). From this perspective, females are rather passive in their mating preference, but males adopting this strategy should theoretically be more likely to sire the majority of the offspring in the mating season, making it more suitable for harem systems or species in which sexual dimorphism prevents females from refusing male mating attempts. When females actively choose their partners, however, it becomes more likely that services serve to increase male’s reputation and hence the probability to be chosen as a mating partner.

Little is known about group movement, organization and progression in vervet monkeys (*Chlorocebus pygerythrus*), compared to baboon species (see for example DeVore 1963; Rhine 1986; Matsumoto-Oda et al. 2018), a primate species with a similar habitat and comparable social organization to vervets. Vervet monkeys live in groups with multiple males and females, where the females exert relatively high levels of mate choice due to female co-dominance (Hemelrijk et al., 2020; Young et al., 2017). Male vervets seem to already show high contributions in some service behaviours before siring any offspring (Baldellou & Henzi, 1992), while female vervets may even “recruit” males during high-risk situations, greeting them more often near rivers where predator encounters are common (Mercier et al., 2017). However, detailed analyses of different male responses to varying predation risk remain scarce.

Here, we test the paternal care and reputation-based partner choice hypotheses (Roberts, 1998; Roberts et al., 2021; van Schaik et al., 2022) in vervet monkeys by examining group progressions in five neighbouring groups. We compared high-risk river crossings – where predator encounters are frequent and terrain is wide and exposed (Mercier et al., 2017) – to low-risk road crossings within the reserve, i.e. crossing of one-track dirt roads that are used by very few slow-driving cars (ours and the owner’s) or not used at all. We predicted that if vervets adjust socio-spatial positioning according to risk, males should disproportionately occupy protective positions at rivers. We expected the dominant male to either lead group progressions, and again more so with increased risk, based on previous studies on primates (dominant male leading foraging movements in chacma baboons: King et al., 2008; dominant male crossing roads first in chimpanzees: Hockings, 2011; Hockings et al., 2006; mountain gorilla males leading the group after risky encounters, Watts, 1994; dominant males found at the front of the group when moving in vervets: Teichroeb et al., 2015, in rhesus macaques: Sueur & Petit, 2008, and in yellow baboons: Rhine & Westlund, 1981). We furthermore expected individuals that are more central in the social network to take up more central physical positions in the group as well (Farine et al., 2017), making them less likely to lead group progressions. If males perform this protective service towards the rest of the group to gain a cooperative reputation and so increase their mating opportunities, we expected males that generally lead risky group progressions to have more mating opportunities in the coming year. Alternatively, if males mainly lead the group during risky situations as a form of paternal care, we expected likely fathers to generally cross first. Threats to infants may differ to some extent from threats to others and thus require incompatible protective strategies. It is therefore possible that dominant males potentially have different strategies based on whether they have sired offspring in the group: dominant males who have not yet sired offspring might be more interested in “showing-off” and ensuring safety of females (who are potential future mates), whereas dominant males who have sired offspring might be more inclined to stay closer to the infants in the group as a protective measure (van Noordwijk & van Schaik, 1988).

If vervets are able to evaluate risk during these progressions and change their behaviour accordingly, this should result in protective socio-spatial organization according to the level of risk, as an adaptive and cooperative response (Hockings et al., 2006). This study will therefore help us to understand the effects of predation risk on group movement behaviour, as well as understand the evolution of cooperative socio-spatial organization.

## Methods

### Behavioural data and progression protocols

Behavioural data were collected at the iNkawu Vervet Project (IVP) field site, Mawana Game Reserve, KwaZulu-Natal, South Africa, from October 2021 until May 2024. Progression behaviour was recorded in five neighbouring groups (AK, BD, KB, LT, NH) of wild-living vervet monkeys (*Chlorocebus pygerythrus*). Vervet monkeys live in promiscuous multimale-multifemale groups with regular male dispersal (Henzi & Lucas, 1980) and philopatric females, who form the social core of the group (Borgeaud et al., 2016; Cheney & Seyfarth, 1990; Seyfarth & Cheney, 2000).

Data are collected year-round by trained field assistants who undergo interobserver reliability and monkey identification tests with the onsite scientific manager. Two protocols were used. From October 2021 to April 2022, only the first and last individuals to cross specific “lines” (roads or rivers) were recorded. From April 2022 onwards, the full crossing order and timing were recorded. Consequently, subgroup structure could only be analysed from April 2022 onwards. Daily group composition was determined from presence matrices, and analyses were restricted to individuals present on the focal date. Only non-natal adult males were considered “males.” Females were classed as adults from age four or when they had offspring; juveniles were <4 years (females) or <5 years (males in natal groups). Our dataset contained no males ≥5 years still in their natal group.

Progressions were defined as crossings of roads or rivers, chosen for visibility and comparable terrain. Roads averaged 5.2 m in width, rivers 43.6 m (width of floodplain). We expected wider, open terrain to correspond to higher risk, and predator encounters are also more common near rivers in our study area (Mercier et al., 2017). We therefore considered rivers the highest-risk terrain. The river that the groups must cross (Hlonyane river) varies in width seasonally, with higher water levels in the summer months (November – March), creating a band of more open terrain due to flooding (Figure 1). While water levels are high, crossing events decrease, and no drowning of individuals has ever been observed. The banks on the edge of the floodplain are generally elevated up to approximately five meters, creating steep cliffs on both sides. Vegetation on these banks is generally high and denser, but between these high banks the lack of trees creates an open landscape. The most common predators of this population of vervets during the day seem to be aerial, with predation attempts and events observed on juvenile and subadult individuals by several species of eagles, but attacks of terrestrial predators, such as caracal, have also been observed as well as predation from snakes such as pythons (unpublished data; EvdW personal observations).

**Figure 1:**
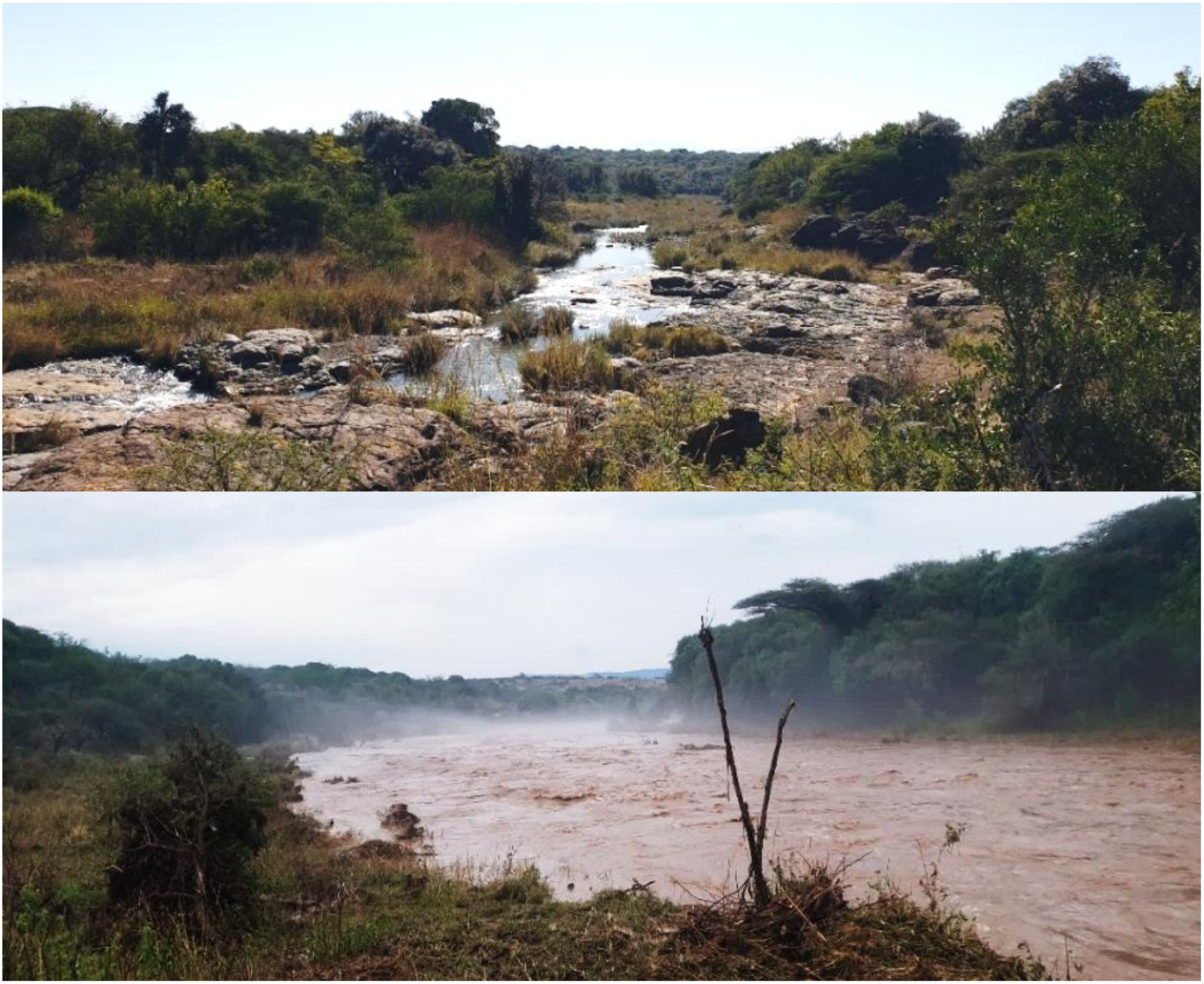
Hlonyane river in the beginning of winter (June, top image) and after heavy rainfall in summer (November, bottom image). Width from riverbank to riverbank can go up to approximately 55 meters.

### Dominance rank, centrality, tenure, fatherhood and habituation calculations

Dominance rank was calculated using the *EloRating* package in R (Neumann & Kulik, 2024; 12 month moving window of conflict data) on each date an individual was recorded in a group progression. The male with the highest score in a group on a given day was classified as “dominant,” all others as “subordinate” (no equal scores ever occurred). Centrality was calculated from grooming networks over the past 12 months using the *socialindices* package (Neumann, 2016). Data were insufficient to calculate reliable rank and centrality for individuals in LT group, who were therefore omitted for this part of the analysis. Scores were standardized relative to the group median (zCSI). Tenure was measured as days since first observation of a male in a non-natal group. Likely fatherhood was inferred from tenure and mating season overlap: males who had not yet experienced a mating season or were present during a mating season (April–July) but before infants were born (Oct–Dec) were considered non-fathers; otherwise, they were likely fathers. Past and future mating opportunities were quantified as the number of mounts recorded in the year preceding and following the progression date. For late 2024, mating data were truncated but still covered the main season when ovulation occurs. Males were classified as “habituated” if born in a study group or present ≥1 year; otherwise they were marked “unhabituated” until reaching one year of tenure.

To ensure adequate sampling of an event, we excluded progressions where identified individuals comprised <10% of those present that day. Because the last individual is only recorded once most of the group has crossed, we did not apply this exclusion to last-crosser tallies. Subgroups were defined by inter-arrival latency: if the time between successive individuals exceeded the global 75th percentile (43 s), the next individual started a new subgroup. The 43-s cut-off is the global 75th percentile of inter-arrival latencies across all events.

### Statistical analyses

All data were analysed in R version 4.2.2. All models were tested to confirm that their assumptions were met and showed robustness in predictive performance.

To assess which age-sex classes led or closed group and subgroup progressions, we ran multinomial models (multinom, *nnet* package; Ripley & Venables, 2016) with age-sex class (adult male, adult female, juvenile) as the response. Four separate models tested leadership of the total group, leadership of subgroups, closing of the total group, and closing of subgroups. Terrain type was the primary predictor, and group identity plus the proportion of males and juveniles in the group were covariates. Proportion of females was omitted to avoid collinearity – as these would always amount to 1 – but alternative coding gave similar results. Unfortunately, the multinomial model did not allow for random effects to be included, which meant we could not account for repeated measures of individuals in these models. Terrain effects were evaluated via planned pairwise contrasts (male vs female; male vs juvenile) within each terrain using *emmeans* on class probabilities (type = “response”). P-values for these limited planned contrasts are reported unadjusted. Data from October 2021 were used for total group leadership/closing, and from April 2022 for subgroup analyses. Leaders of the total group were excluded from subgroup-leader models, and single-individual subgroups were excluded from subgroup-closer models. Results on leading and closing subgroups can be found in the supplementary results.

A binomial GLMM tested whether males were more likely to cross alone when leading and whether this varied by terrain. Data from April 2022 onwards were used. For this analysis, we only used males that were recorded during crossings in which a male was recorded as leading the group. The response variable was whether a male crossed with others (y/n). Predictors were terrain type, leadership status, and their interaction, with group identity as a control. Random intercepts were included for individual and date.

A second binomial GLMM investigated factors predicting which male tended to lead a progression. Data from October 2021 onwards was used for this model. Each observation represented a male present during a male-led event. The response was whether that male led (1) or not (0). The log of the number of males present was included as an offset to account for variation in male availability. Predictors were terrain type, dominance status, centrality, potential fatherhood, past and future mating success, and habituation. Because fatherhood and habituation are tenure-based, tenure itself was not included in the final model. A model with tenure as a predictor instead of potential fatherhood and habituation was also performed but performed less well and did not show any effects of tenure itself. Group identity and season were included as fixed effects to control for group-specific and seasonal variation. Random intercepts were included for individual ID. LT group was omitted due to insufficient data for rank/centrality. We also tested interactions between dominance and fatherhood, and between dominance and both mating measures, given strong effects of dominance.

A final GLMM (negative binomial; *glmmTMB;* Brooks et al., 2017) tested whether non-leader males crossed in subgroups containing infants. Data from April 2022 onwards in four groups (excluding LT) were used. The response variable was the count of infants in the subgroup of a non-leader male. Predictors were terrain type in interaction with dominance status, fatherhood, and centrality. Habituation and season were again included as controls. Group and male ID were random effects, with number of males in the group as an offset. Because data for this analysis started in April 2022, the dominant non-father category lacked sufficient instances of crossing with infants (likely because these males generally led), the dominance- fatherhood interaction could not be estimated; we therefore ran a complementary Firth logistic regression with a binary response (crossed with infant: yes/no).

For an overview of all the models performed, see Table S2.

## Results

We had a total of 406 observations on group progression in which the first individual to cross was recorded between October 2021 and May 2024. Across these 406 group-progression events (177 roads; 229 rivers), males led 52.7% overall. When the group crossed a road, males generally led 45.0 % of progressions, whereas they led 58.6 % of group progressions when crossing rivers. We had 260 observations during the same period on group progression in which the last individual to cross was recorded. Of those observations, 110 concerned road crossings and 150 river crossings. Males were the last to cross in 31.9 % of observations. Males closed group progression in 23.6 % of roads progressions and 38.0 % of river progressions. Males constituted a median of 16.2% of group membership across group-days (Table 1).

**Table 1:** group composition and size, number of crossings recorded per group.

| Group | Prop. males<br>(median) | Group size<br>(median) | Number of observations on leaders of progression over |  | Number of observations on closers of progression over |  |
| --- | --- | --- | --- | --- | --- | --- |
|  |  |  | Roads | Rivers | Roads | Rivers |
| AK | 0.15 | 23 | 80 | 0 | 63 | 0 |
| BD | 0.21 | 68 | 22 | 77 | 9 | 51 |
| KB | 0.18 | 24 | 35 | 11 | 3 | 26 |
| LT | 0.09 | 31 | 0 | 21 | 0 | 22 |
| NH | 0.18 | 46 | 44 | 70 | 35 | 51 |

Adult males were generally more likely to lead group progressions – no matter the terrain – than juveniles (*estimate* = 0.36, *SE* = 0.05, *p* < 0.001) and adult females (*estimate* = 0.26, *SE* = 0.05, *p* = 0.0003). We found an effect of terrain on the probability of a male to lead (Figure 2): males lead the group during progressions over roads nearly significantly more than females (*estimate* = 0.19, *SE* = 0.08, *p* = 0.09) and juveniles (*estimate* = 0.21, *SE* = 0.08, *p* = 0.06), but did lead group progressions over rivers significantly more than females (*estimate* = 0.34, *SE* = 0.07, *p* = 0.0003) and juveniles (*estimate* = 0.51, *SE* = 0.05, *p* < 0.0001). The probability of a male to lead increased significantly as well over rivers compared to roads (*estimate* = 0.15, *SE* = 0.06, *p* = 0.03). The group identity did not influence the probability of an age-sex class leading (*Chi^2^ =* 12.16, *df =* 8, *p* = 0.14).

**Figure 2:**
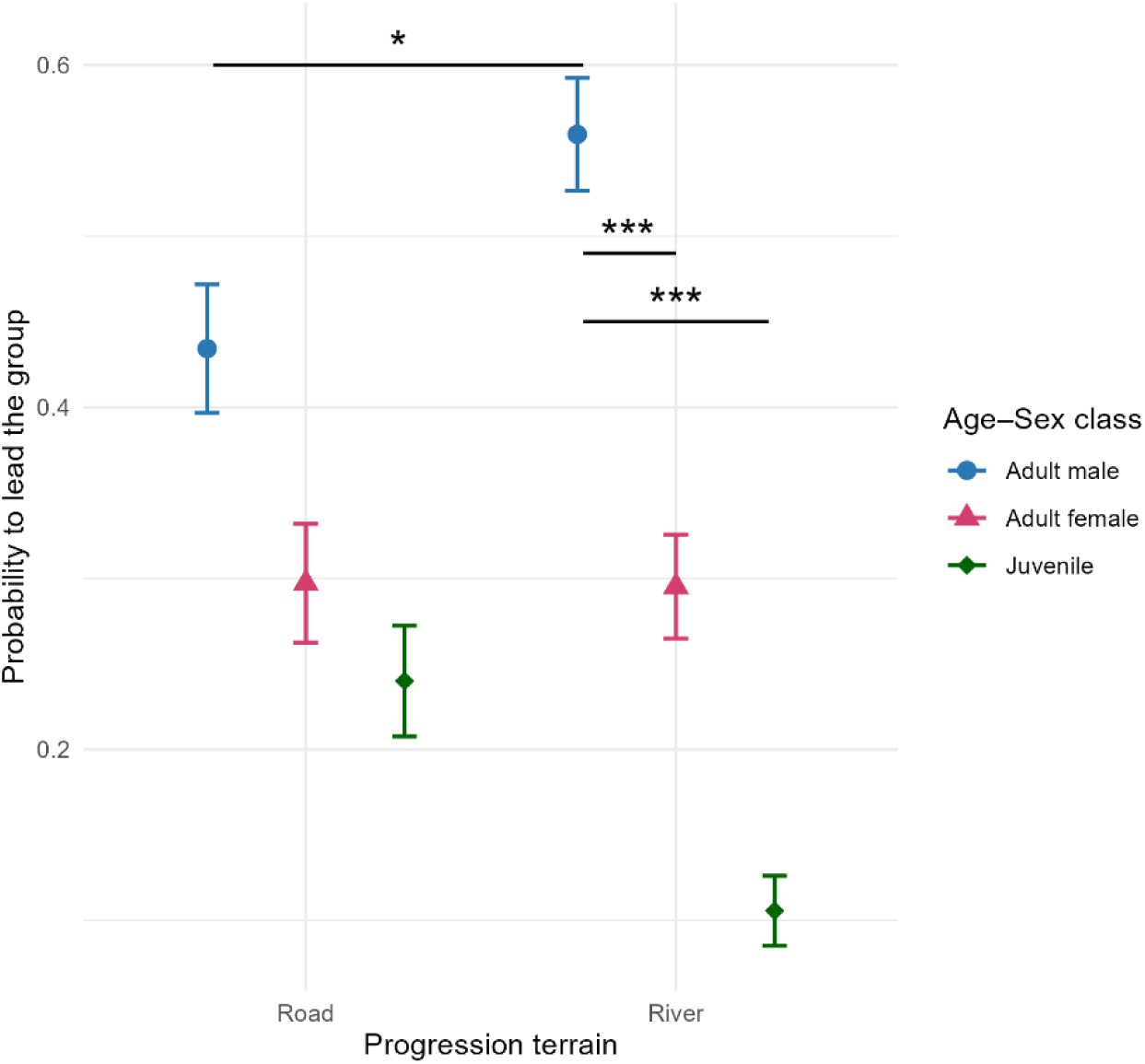
The probability of leading the group (y-axis) over different progression terrains (x-axis), per age and sex classes. Males were significantly more likely to lead in river progressions than adult females and juveniles (indicated with ***), and the probability of a male leading was higher in river progressions than in road progressions (indicated with *).

Males were not more or less likely to cross last than females (*estimate* = 0.09, *SE* = 0.06, *p* = 0.31) or juveniles (*estimate* = 0.02, *SE* = 0.07, *p* = 0.95) irrespective of the terrain, or in either terrain types (roads – male vs female: *estimate* = -0.07, *SE* = 0.11, *p* = 0.77; roads – male vs juvenile: *estimate* = -0.06, *SE* = 0.12, *p* = 0.83; rivers – male vs female: *estimate* = -0.12, *SE* = 0.08, *p* = 0.37; rivers – male vs juvenile: *estimate* = 0.11, *SE* = 0.07, *p* = 0.25). The probability of a male to close a progression did not differ over terrain types either (*estimate* = -0.05, *SE* = 0.08, *p* = 0.58). Females were more likely to cross last during river progressions than juveniles (*estimate =* 0.23, *SE =* 0.07, *p =* 0.01), but not during road progressions (*estimate =* 0.007, *SE* = 0.12, *p =* 0.99). The group identity marginally non-significantly affected the probability of an age-sex class closing (*Chi^2^* = 14.28, *df =* 8, *p =* 0.07), where in AK group adult females are more likely to close than adult males (*estimate* = 0.40, *SE* = 0.14, *p* = 0.03) and juveniles (*estimate* = 0.06, *SE* = 0.09, *p* = 0.81). It is worth noting here that no river progressions were observed in AK group (see Table 1).

We found that when males lead the group, they were followed on average by other individuals within 43 seconds (the statistical result to define membership to the same subunit) in 33.7 % of observed group progressions (see Figure 3). Males that did not lead the group generally crossed within 43 seconds of other individuals in 80.9 % of observations, which was significantly different from males leading (*estimate* = 2.24, *SE* = 0.34, *p* < 0.0001). We also analysed the two crossing contexts separately. When crossing roads, males that did not lead crossed close to others 82.9 % of the time, while the leader crossed close to others for 51.2 % of the time (*estimate* = 1.49, *SE* = 0.49, *p* = 0.002). When crossing rivers, males that did not lead crossed close to others 80.3 % of the time, and males that led crossed close to others 20.0 % of the time (*estimate* = 2.24, *SE* = 0.34, *p* < 0.0001). Males were more likely to lead alone in rivers than in roads (*estimate* = 2.10, *SE* = 0.63, *p* = 0.0008) but not more or less likely to cross with others when not leading in rivers compared to roads (*estimate* = 0.60, *SE* = 0.52, *p* = 0.25).

**Figure 3:**
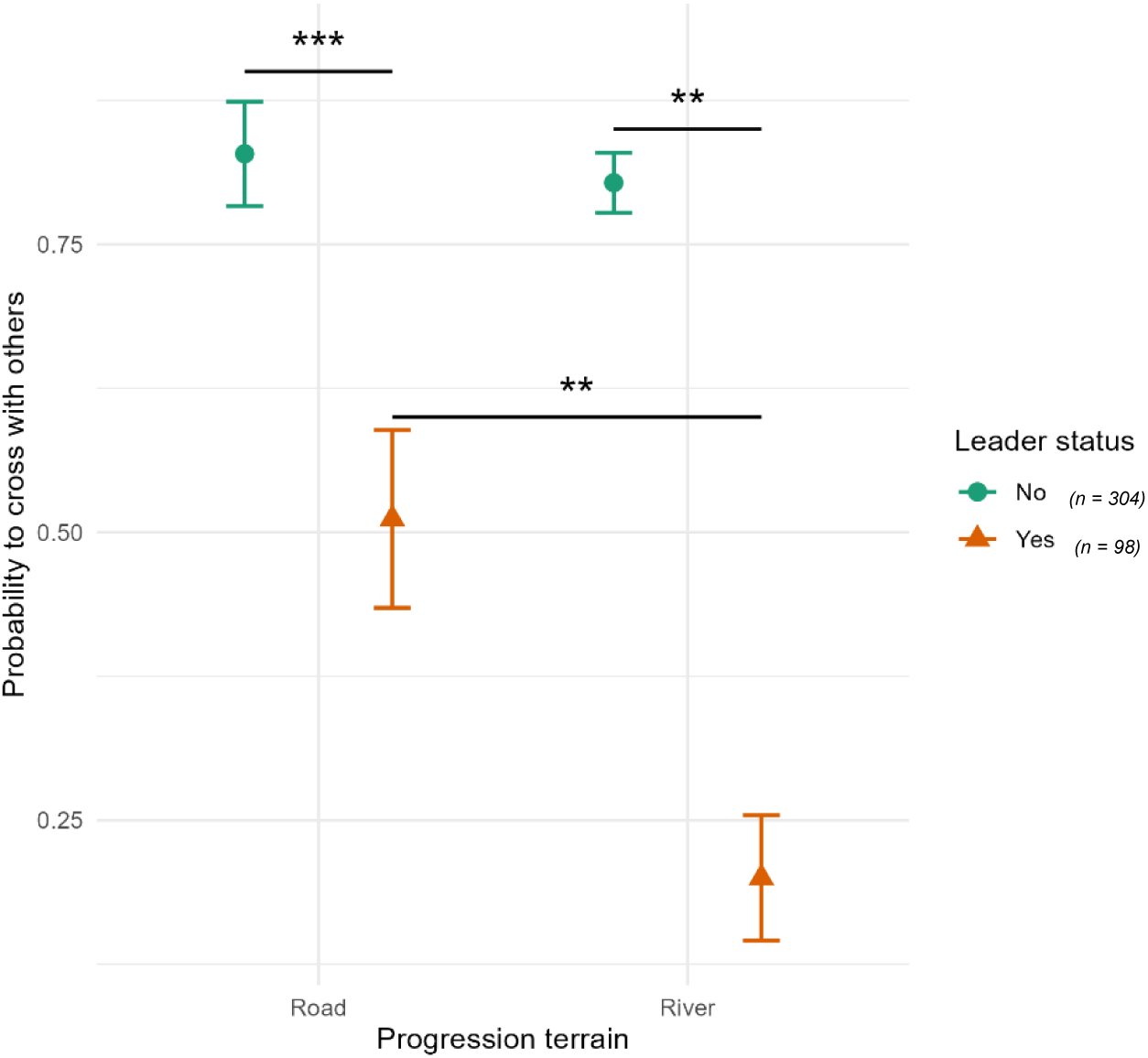
The probability of males to cross in close proximity of others (y-axis) over different progression terrains (x-axis), per whether a male was leading (yes or no). Males were significantly more likely to cross alone when leading in both river and road progressions (indicated with ***), and the probability of a male leading alone was higher in river progressions than in road progressions (indicated with **).

Dominant males were more likely to lead group progressions than subordinate males during river crossings (*estimate* = 2.31, *SE* = 0.67, *p* = 0.0005), but not during road crossings (*estimate* = 0.57, *SE* = 0.62, *p* = 0.36). Furthermore, the probability of the dominant male leading the group was higher in river progressions than in road progressions (*estimate* = 1.60, *SE* = 0.81, *p* = 0.05). Additionally, dominant non-fathers were marginally non-significantly more likely to lead in river progressions than in road progressions (*estimate* = 2.57, *SE* = 1.49, *p* = 0.09), marginally non-significantly more likely to lead in river progressions than dominant fathers (*estimate* = 2.28, *SE* = 1.27, *p* = 0.07), more likely to lead group progressions in rivers than subordinate non-fathers (*estimate* = 3.97, *SE* = 1.29, *p* = 0.002) and subordinate fathers (*estimate* = 2.93, *SE* = 1.25, *p* = 0.02; Figure 4). Dominant fathers were more likely to lead group progressions over rivers than subordinate fathers (*estimate* = 0.66, *SE* = 0.31, *p* = 0.03) and subordinate non-fathers (*estimate* = 1.69, *SE* = 0.52, *p* = 0.001). Subordinate fathers were more likely to lead during river progressions than subordinate non-fathers (*estimate* = 1.03, *SE* = 0.46, *p* = 0.02). We found no differences in the likelihood to lead between dominant or subordinate fathers or non-fathers in road progressions. Based on a priori hypotheses regarding the role of fatherhood in dominant males, we report the unadjusted p-values for these key contrasts. These results are consistent with the raw data patterns observed (Figure 4).

**Figure 4:**
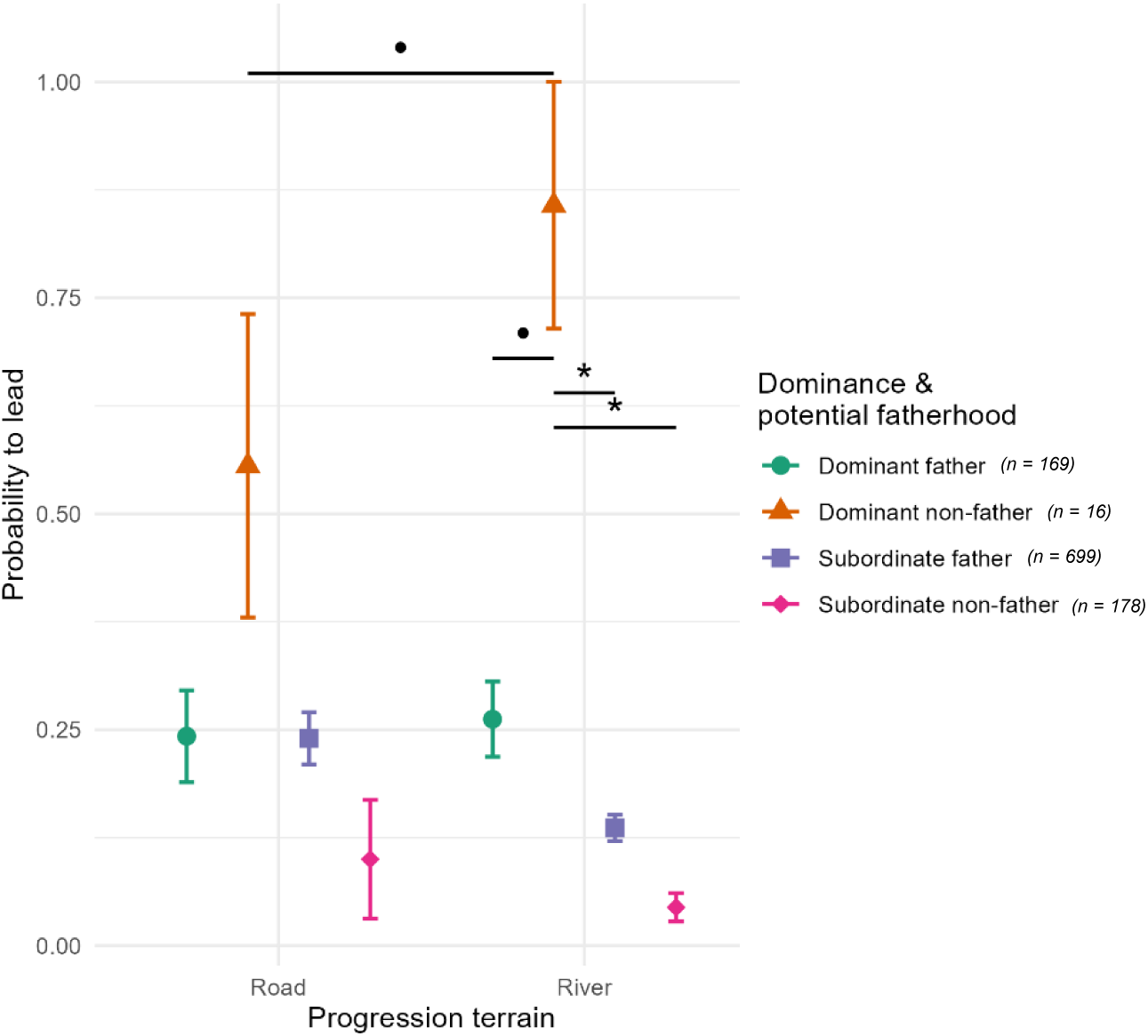
The probability of males to lead group progression (y-axis) over different terrain (x-axis), as a function of dominance and potential fatherhood status. Dominant non-fathers (orange triangles) are more likely to lead group progressions over rivers than over roads, and more likely than all other male categories within river progressions.

We found no effect of centrality on the probability to lead group progressions over roads (*estimate* = -0.22, *SE* = 0.19, *p* = 0.24) or rivers (*estimate* = -0.19, *SE* = 0.15, *p* = 0.21). The mating opportunities of a male in the twelve months prior to the crossing event did not influence his probability to lead the group in roads (*estimate* = -0.58, *SE* = 0.40, *p* = 0.15) or rivers (*estimate* = -0.11, *SE* = 0.19, *p* = 0.54). There was a negative trend between past mating opportunities and a dominant male’s probability to lead over roads (*estimate* = -1.32, *SE* = 0.75, *p* = 0.08), but this trend did not differ significantly from the other non-significant slopes (difference between slopes: *Chi^2^ =* 2.22, *df* = 1, *p* = 0.14; dominant male’s probability to lead over rivers: *estimate* = -0.23, *SE* = 0.33, *p* = 0.48; subordinate male’s probability to lead over roads: *estimate* = 0.16, *SE* = 0.22, *p* = 0.47; subordinate male’s probability to lead over rivers: *estimate* = 0.008, *SE* = 0.15, *p* = 0.96). Although we did not find a difference in the effect of dominance, future mating opportunities and terrain on the type of a male to lead (*Chi^2^*= 0.19, *df* = 1, *p* = 0.66), we found a negative trend between future mating success and probability to lead in river progressions (*estimate* = -0.33, *SE* = 0.19, *p* = 0.08). This seemed especially so for dominant males, as we found a negative trend between future mating success and probability to lead over rivers for dominant males (*estimate* = -0.56, *SE* = 0.33, *p* = 0.09), but not for subordinate males (*estimate* = -0.09, *SE* = 0.14, *p* = 0.54). However, as these slopes did not significantly differ from each other, these results should be interpreted with caution. We found no significant or meaningful effects of our other parameters (season, habituation and group identity). Although group identity did show significant effects, this was due to the number of males in each group – and therefore not meaningful for our results.

Our model testing whether males that do not lead cross more often with infants when they are potential fathers yielded negative results: the number of infants in a male’s subgroup were not affected by his dominance status (*Chi^2^* = 0.03, *df* = 1, *p* = 0.87) as well as the fatherhood status (*Chi^2^* = 0.16, *df* = 1, *p* = 0.67). Instead, a male’s centrality predicted the number of infants in his subgroup, positively (*Chi^2^* = 5.79, *df* = 1, *p* = 0.01). This seemed to be mainly driven by the positive slope of centrality during river progressions (*estimate* = 0.33, *SE* = 0.13, *p* = 0.01). A male’s centrality did not predict the number of infants in his subgroup during road progressions (*estimate* = 0.13, *SE* = 0.33, *p* = 0.68). Our Firth’s model dealing with complete separation included the interaction between dominance and fatherhood but yielded similar results: whether a male crossed with an infant in his subgroup was unaffected by his dominance status (*Chi^2^* = 0.02, *df* = 1, *p* = 0.47), his paternal status (*Chi^2^* = 0.01, *df* = 1, *p* = 0.93) or an interaction between the two (*Chi^2^* = 0.03, *df* = 2, *p* = 0.86).

Males did not seem to take on any specific roles within subgroups, neither leading nor closing them. For detailed results on subgroup progressions, see the supplementary results.

## Discussion

This study compared the spatial positioning of vervet monkeys during high- and low-risk situations, to test whether the male bias in antipredator behaviour reflects parental care or reputation-based mate choice (Roberts, 1998; Roberts et al., 2021; van Schaik et al., 2022). We found that adult males were more likely to assume leadership roles during progressions over rivers – terrain characterized by higher exposure and predation risk – than across low-risk terrains. In these progressions, males positioned themselves at the front, well ahead of the group, thereby increasing their individual predation risk. Dominant males were most likely to lead, particularly those without offspring, while we found no convincing effect of centrality or mating opportunities on the likelihood of leadership. These findings suggest that the pattern of male leaders traversing risky terrain alone is not merely a by-product of group movement dynamics but a protective and thus, cooperative service.

Our findings that males often lead alone during high risk progressions support the protection hypothesis, which posits that strong individuals strategically position themselves in risky locations to safeguard the group (Rhine et al., 1981; Rhine & Westlund, 1981), by increasing their own individual predation risk due to physical isolation (Romenskyy et al., 2020; Suire et al., 2023). Such isolated leadership appears to be a deliberate strategy undertaken predominantly by dominant males. These males, potentially stronger and with higher status, may take on riskier positions to draw attention of predators and/or scout unexplored terrain ahead. This is in line with the male-bias that was found in anti-predatory services described by van Schaik and colleagues (2022) and suggests that vervet monkeys are able to adapt to differing risks with adaptive and cooperative socio-spatial organisation. However, other strategic positions, such as the rear, were not necessarily occupied by males. One plausible interpretation for this lack of males in the rear is that closing the group in this species is not necessarily particularly risky, as the individuals at the rear may benefit from the protection afforded by those ahead of them. Although this is not directly in line with the protection hypothesis (Rhine et al., 1981; Rhine & Westlund, 1981), from which it can be hypothesized that individuals occupying the group’s edges – therefore including the rear – are always more vulnerable, it is in line with the general trend found in van Schaik et al. (2022). Another explanation for this lack of males closing group progressions might be that the crossing itself is not the most dangerous, but whatever lies ahead on the other side. If that is the case, individuals that go ahead of the group – especially doing so alone – are the only ones able to perform a service: crossing with others or close to infants has no “effect” if the real danger does not lie in the crossing itself.

When males did not lead, they usually crossed with other individuals (i.e., within subgroups) and were unlikely to take the front or rear of these subgroups. Importantly, males were not more likely to cross with infants based on paternal status. While we expected fathers to cross with potential offspring, we found no such effect, refuting the paternal care hypothesis. Males therefore do not appear to provide protective services to specific offspring, either by leading or staying close. Nevertheless, their general tendency to cross with others may indicate investment in female survival and future mating opportunities (Bshary et al., 2022), if the progression itself is perceived as dangerous. However, when taken together, these results suggest that the main service during group progression is likely scouting ahead rather than remaining close to vulnerable group members.

It could be argued that male leadership during risky progressions is a by-product of having high dominance status, consistent with previous studies (Hockings et al., 2006; Hockings, 2011; King et al., 2008; Rhine & Westlund, 1981; Sueur & Petit, 2008; Teichroeb et al., 2015; Watts, 1994). Likewise, proximate accounts such as greater physical strength and caloric needs (potentially favouring early access to resources; King et al., 2008), higher boldness or confidence (Petit & Bon, 2010; Hockings et al., 2006; Rhine & Westlund, 1981), and a higher probability of being followed at departure (Sueur & Petit, 2008) could increase baseline leadership among dominant males in general. However, these accounts do not on their own predict the risk-contingent increase we observed: dominant males specifically became more likely to lead as risk increased. Moreover, dominant males can typically monopolise food resources, reducing the necessity of reaching food sources first. Thus, while such mechanisms may facilitate leadership, they are insufficient to explain why leadership by dominants intensifies under high risk.

According to the reputation-based partner choice hypothesis, dominant males, who are potentially more likely to sire future offspring in the group, may demonstrate value by leading in risky contexts and thereby improving female survival and gain increased reproductive opportunities. Nonetheless, we did not find mating advantages for males that lead, and even a negative trend for our dominant males. One explanation may be our use of mating rather than paternity success, since female primates often mate strategically (Fujita, 2010; Inoue & Takenaka, 2008). Another could be group size: the unusually large groups in our study may limit monopolization and increase female choice, potentially explaining the lack of expected mating advantages (Freeman et al., 2016; Hemelrijk et al., 2020). Evaluation of long-term data on mating and paternity is needed to resolve this issue.

Nevertheless, the lack of evidence of reputation-based partner choice – especially when it comes to mating advantages – as well as the clear differentiation between paternal and dominance status warrants that we consider other potential explanations for this particular male service. One possibility is that dominant males, especially newly immigrated non-fathers, lead progression in dangerous terrain as a signal of quality towards other males. If leading stabilises a male’s own rank, it yields benefits due to, for example, continued priority of access to resources. In Arabian babblers for example, subordinate males might mob predators to signal quality to other likely-to-disperse males (Maklakov, 2002), while dominant males were found to interfere and replace subordinate males from guarding positions, potentially competing over performing cooperative acts (Dattner et al., 2016). These examples illustrate that cooperative acts can function as signals in between male competition, not only male-female interactions. Applying this to vervets, dominant non-fathers may lead risky crossings not to attract mates but to assert or stabilise their rank relative to other males. Taken together, these patterns point to complementary male strategies: dominant non-fathers assume high-risk leadership roles that may function both as group protection and as a social signal, whereas other males contribute by travelling with the group without preferentially being close to vulnerable individuals. This interpretation highlights that dominance is necessary but not sufficient for risky leadership; instead, its expression depends jointly on ecological risk and a male’s reproductive and social context.

Overall, our findings suggest that male leadership in vervet monkeys is shaped by a combination of group-protective behaviour and potential signalling functions within male hierarchies. While our findings support the interpretation that male leadership in risky contexts is a deliberate cooperative service, we found no evidence that it translates into mating advantages. Furthermore, dominant males preferentially assumed high-risk positions, likely as a protective service to the group, thereby adapting their behaviour based on potential presence of offspring, changing their cooperative behaviour according to life-history payoffs. These insights highlight the role of risk management and service provisioning in shaping primate social organisation, while also calling for further research into whether protective service provisioning confers evolutionary benefits in terms of reproductive success. Future work incorporating paternity data, long-term rank trajectories, and experimental risk manipulation will be essential to disentangle the protective and signalling components of this male service.

## Acknowledgements

We want to thank the team at IVP for their crucial contributions to data collection, group monitoring and help with logistics, and to everyone who got their feet wet trying to record a crossing: thank you. We want to thank Nokubonga Dhlamini, Michael Henshall, Siboniso Thela and Zonke Mbutho for running the field site, as well as supporting the field team. We are grateful to the van der Walt family for permission to work in their reserve, as well as to Ezemvelo KZN Wildlife for granting us the necessary permissions to do so. Radu Slobodeauanu provided statistical assistance, for which we are thankful. We thank Maria Granell Ruiz for her helpful input on the interpretation of the results and the valuable discussions. This research was supported by the Swiss National Science Foundation (grant number 310030_197884). The field costs during data collection were funded by grants to EvdW from the Swiss national Science Foundation (grant number PP00P3_198913), the grant ‘ProFemmes’ of the Faculty of Biology and Medicine of the University of Lausanne and the European Research Council under the European Union’s Horizon 2020 research and innovation program for the ERC ‘KNOWLEDGE MOVES’ starting grant (grant agreement No. 949379).

For the purpose of Open Access, a CC BY public copyright license is applied to any Author Accepted Manuscript (AAM) version arising from this submission.

## Supplementary results

Males that did not lead the group were less likely to lead subgroups irrespective of terrain than females (*estimate* = 0.26, *SE* = 0.02, *p* < 0.0001) and juveniles (*estimate* = 0.21, *SE* = 0.02, *p* < 0.0001). These differences did not differ by crossing type either (roads – male vs. female: *estimate* = -0.28, *SE* = 0.02, *p* < 0.0001; roads – male vs. juvenile: *estimate* = -0.27, *SE* = 0.02, *p* < 0.0001; rivers – male vs. female: *estimate* = -0.26, *SE* = 0.02, *p* < 0.0001; rivers – male vs. juvenile: *estimate* = -0.15, *SE* = 0.02, *p* < 0.0001). The probability of a male to lead a subgroup differed significantly between the crossing types (road – river: *estimate* = -0.05, *SE* = 0.02, *p* = 0.006). Females were slightly more likely to lead subgroups than juveniles irrespective of terrain (*estimate* = 0.06, *SE* = 0.03, *p* = 0.08), which was mainly explained by an increased likelihood of females leading subgroups during river progressions than juveniles (*estimate* = 0.11, *SE* = 0.03, *p* = 0.006). Females were not more likely to lead subgroups during road progressions than juveniles (*estimate* = 0.01, *SE* = 0.04, *p* = 0.93) and their likelihood of leading did not significantly differ between roads and rivers (*estimate* = -0.02, *SE* = 0.02, *p* = 0.30).

Males were also less likely to close subgroups, irrespective of the terrain, than females (*estimate* = -0.13, *SE* = 0.02, *p* < 0.0001) and juveniles (*estimate* = -0.29, *SE* = 0.02, *p* < 0.0001). Males were less likely to close subgroups in both road and river crossings than both females (roads: *estimate* = -0.15, *SE* = 0.02, *p* < 0.0001; rivers: *estimate* = -0.11, *SE* = 0.02, *p* = 0.0002) and juveniles (roads: *estimate* = -0.32, *SE* = 0.03, *p* < 0.0001; rivers: *estimate* = -0.26, *SE* = 0.02, *p* < 0.0001). The probability of a male to close a subgroup was non-significantly higher in rivers than in roads (*estimate* = 0.03, *SE* = 0.02, *p* = 0.06). Females were less likely to close subgroups than juveniles irrespective of terrain (*estimate* = -0.16, *SE* = 0.02, *p* < 0.0001), which was consistent in both roads (*estimate* = -0.17, *SE* = 0.03, *p* = 0.0001) and rivers (*estimate* = -0.15, *SE* = 0.03, *p* = 0.0001). The probability of closing a subgroup remained consistent between roads and rivers for both females (*estimate* = 0.003, *SE* = 0.02, *p* = 0.88) and juveniles (*estimate* = 0.03, *SE* = 0.02, *p* = 0.18).

When considering the probability to lead during different life-stages of individual males that were recorded as dominant non-fathers, we found that their trajectories were similar to the static pattern among all males (Figure 5): these males were most likely to lead river progressions when they were dominant non-fathers (83.3 %), and less when they were subordinate fathers (54.5 %), dominant fathers (30.8 %) and especially subordinate non-fathers (25.0 %). This simplified model yielded similar results: the paternal and dominance status of the males significantly affected their probability to lead (*Chi^2^* = 9.16, *df* = 3, *p* = 0.03). Dominant non-fathers were almost significantly more likely to lead than when they were dominant fathers (*estimate* = 4.18, *SE* = 1.75, *p* = 0.08), although not more likely than when they were subordinate fathers or non-fathers (resp.: *estimate* = 2.55, *SE* = 1.58, *p* = 0.37; *estimate* = 2.52, *SE* = 1.60, *p* = 0.39), although this might be attributed to lacking statistical power due to our small sample size (n = 6 dominant non-father; 13 dominant father; 11 subordinate father; 4 subordinate non-father). We found slight seasonal effects as well (*Chi^2^*= 5.43, *df* = 2, *p* = 0.07), where these males were marginally non-significantly more likely to lead group progression over rivers during the summer than during the baby season (*estimate* = 3.86, *SE* = 1.86, *p* = 0.09). Other seasons did not differ, but data was lacking in the mating season.

**Figure 5:**
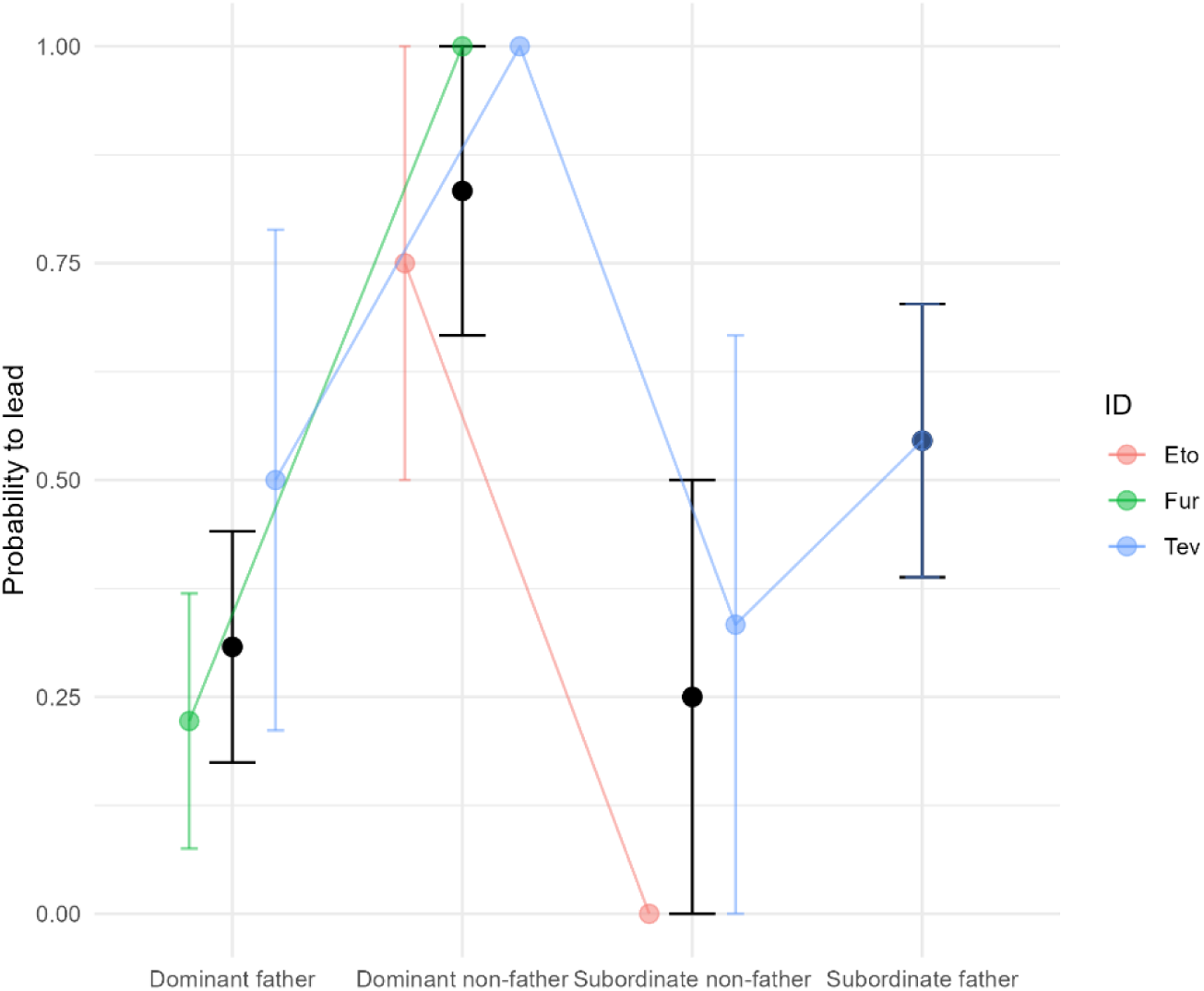
The probability of the males that were recorded in river progressions as dominant non-fathers to lead group progression (y-axis) over different dominance and paternity statuses (x-axis). The colours represent the three different males.

## Supplementary materials

**Table S1:** overview of all males used in this study. First column *Male* gives the male’s ID, second *Origin* column shows where the male was born, *Group* indicates the focal group in which progression data has been collected. *Tenure @ first (d)* gives the tenure in days on the first day the male was recorded in a progression. *Hab @ first* indicates whether the male was habituated (H) or not (U) when data was first recorded. *Father @ first* gives whether the male was a potential father on the first day data was recorded. *zCSI (mean/med)* and *Elo (mean/med)* give the zCSI and Elo score (centrality and rank) as calculated using the method described in the methods section. It gives both the mean and the median score over all observations. *Infants (mean)* gives the average number of infants in this male’s subgroup while crossing. *Mounts -12m* and *Mounts +12m* give the average recorded number of mounts during the previous (*-12m*) and coming (*+12m*) year over all observations. *# GP leads* indicates the number of times the male was recorded leading the group, *# SubGP leads* the number the male was recorded leading a subgroup.

| Male | Origin | Group | Tenure<br>@ first<br>(d) | Hab<br>@ first | Father<br>@ first | zCSI<br>(mean/med) | Elo<br>(mean/med) | Infants<br>(mean) | Mounts<br>-12m | Mounts<br>+12m | # GP<br>leads | # SubGP<br>leads |
| --- | --- | --- | --- | --- | --- | --- | --- | --- | --- | --- | --- | --- |
| Apa | Unknown/<br>Non-study | BD | 0 | U | No | -1.16 (-1.42) | 950 (962) | 0.50 | 0.50 | 1.00 | 0 | 2 |
| Boo | Unknown/<br>Non-study | BD | 12 | U | No | -1.25 (-1.34) | 1072 (1050) | 1.83 | 2.33 | 23.17 | 0 | 4 |
| Bra | Unknown/<br>Non-study | BD | 19 | U | No | -1.34 (-1.36) | 1106 (1034) | 0.05 | 4.00 | 34.19 | 8 | 14 |
| Buk | Unknown/<br>Non-study | AK | 175 | U | No | -0.86 (-1.10) | 1059 (1071) | 0.07 | 21.45 | 21.19 | 19 | 29 |
| Cai | Unknown/<br>Non-study | NH | 9 | U | No | 0.92 (1.89) | 1377 (1377) | 0.00 | 3.67 | 16.33 | 1 | 1 |
| Dak | Unknown/<br>Non-study | NH | 11 | U | No | -1.23 (-1.21) | 1011 (1020) | 0.68 | 0.91 | 15.09 | 0 | 2 |
| Dal | LT | BD | 11 | H | No | -0.98 (-1.06) | 806 (856) | 1.61 | 8.56 | 20.72 | 0 | 6 |
| Dix | LT | BD | 11 | H | No | -1.29 (-1.34) | 778 (731) | 1.22 | 7.83 | 29.11 | 0 | 2 |
| Dok | CR | BD | 879 | U | Yes | -0.25 (-0.01) | 838 (818) | 0.05 | 12.45 | 10.95 | 2 | 2 |
| Dri | Unknown/<br>Non-study | BD | 2 | U | No | -0.08 (-0.94) | 824 (868) | 1.35 | 1.76 | 13.71 | 2 | 2 |
| Eto | Unknown/<br>Non-study | KB | 143 | U | No | -1.70 (-1.70) | 1088 (1100) | 0.00 | 4.00 | 0.00 | 3 | 3 |
| Fle | Unknown/<br>Non-study | NH | 451 | U | Yes | -0.42 (-0.94) | 874 (869) | 0.87 | 3.69 | 3.50 | 7 | 23 |
| Flu | Unknown/<br>Non-study | BD | 909 | U | Yes | -0.42 (-0.46) | 1060 (1051) | 0.25 | 12.52 | 7.73 | 5 | 11 |
| Fur | Unknown/<br>Non-study | KB | 5 | U | No | -0.28 (-0.41) | 1036 (1025) | 1.43 | 5.49 | 2.06 | 13 | 37 |
| Hav | Unknown/<br>Non-study | NH | 9 | U | No | -1.29 (-1.33) | 1134 (1134) | 0.33 | 1.83 | 20.42 | 0 | 0 |
| Kno | Unknown/<br>Non-study | BD | 2 | U | No | -1.38 (-1.38) | 787 (807) | 0.62 | 6.43 | 2.90 | 4 | 14 |
| Kny | Unknown/<br>Non-study | NH | 10 | U | No | -1.21 (-1.24) | 883 (890) | 0.23 | 5.56 | 29.15 | 5 | 7 |
| Kom | Unknown/<br>Non-study | BD | 713 | U | Yes | -1.13 (-1.11) | 1048 (1084) | 0.66 | 4.24 | 2.98 | 13 | 23 |
| Lif | Unknown/<br>Non-study | NH | 146 | U | No | -1.11 (-1.09) | 1122 (1061) | 0.58 | 16.84 | 31.57 | 12 | 14 |
| Mat | AK | BD | 616 | H | Yes | -0.02 (-0.01) | 699 (699) | 0.00 | 26.00 | 3.00 | 0 | 0 |
| Nak | AK | BD | 34 | H | No | -0.68 (-0.70) | 950 (901) | 1.76 | 18.36 | 10.78 | 2 | 6 |
| Nge | AK | BD | 7 | H | No | -1.03 (-0.96) | 827 (814) | 0.97 | 18.21 | 10.77 | 1 | 5 |
| Oke | Unknown/<br>Non-study | KB | 145 | U | No | 0.46 (0.30) | 906 (912) | 0.29 | 11.41 | 5.59 | 4 | 6 |
| Pal | CR | BD | 1,783 | U | Yes | -0.22 (-0.21) | 1322 (1318) | 0.00 | 9.44 | 0.00 | 0 | 0 |

| Male | Origin | Group | Tenure<br>@ first<br>(d) | Hab<br>@<br>first | Father<br>@ first | zCSI<br>(mean/med) | Elo<br>(mean/med) | Infants<br>(mean) | Mounts<br>-12m | Mounts<br>+12m | # GP<br>leads | # SubGP<br>leads |
| --- | --- | --- | --- | --- | --- | --- | --- | --- | --- | --- | --- | --- |
| Pom | BD | NH | 1 | H | No | -1.07 (-1.05) | 904 (908) | 0.00 | 13.44 | 41.50 | 10 | 10 |
| Pro | NH | BD | 835 | H | Yes | 0.42 (0.70) | 1655 (1660) | 0.00 | 40.32 | 1.16 | 8 | 8 |
| Que | Unknown/<br>Non-study | KB | 33 | U | No | -1.15 (-1.09) | 923 (942) | 0.48 | 15.41 | 15.52 | 4 | 8 |
| Sey | CR | BD | 888 | U | Yes | -0.76 (-0.77) | 1072 (1102) | 1.17 | 15.71 | 15.23 | 9 | 25 |
| Sho | Unknown/<br>Non-study | AK | 385 | U | Yes | -0.97 (-0.96) | 950 (934) | 0.30 | 10.30 | 13.00 | 16 | 24 |
| Sio | Unknown/<br>Non-study | NH | 283 | U | Yes | -0.09 (-0.10) | 696 (701) | 0.00 | 15.00 | 0.00 | 1 | 1 |
| Tam | Unknown/<br>Non-study | NH | 528 | U | Yes | -1.06 (-1.10) | 922 (917) | 0.27 | 4.22 | 1.73 | 38 | 42 |
| Tch | Unknown/<br>Non-study | AK | 175 | U | No | 0.37 (0.37) | 941 (941) | 0.00 | 6.83 | 12.85 | 1 | 1 |
| Tch | Unknown/<br>Non-study | NH | 4 | U | No | -1.23 (-1.26) | 941 (941) | 0.61 | 6.83 | 12.85 | 8 | 12 |
| Ted | CR | BD | 884 | U | Yes | -1.20 (-1.15) | 688 (675) | 0.00 | 11.05 | 0.58 | 1 | 1 |
| Tev | Unknown/<br>Non-study | KB | 145 | U | No | -0.58 (-0.58) | 993 (988) | 0.32 | 15.63 | 10.73 | 20 | 26 |
| Tow | Unknown/<br>Non-study | BD | 37 | U | No | -1.04 (-1.02) | 981 (981) | 0.00 | 1.25 | 16.75 | 6 | 6 |
| Umb | CR | BD | 882 | U | Yes | -0.74 (-0.94) | 1110 (1093) | 0.45 | 13.76 | 8.41 | 7 | 35 |
| Utr | Unknown/<br>Non-study | NH | 510 | U | Yes | -0.92 (-0.92) | 994 (975) | 0.18 | 10.27 | 8.84 | 14 | 48 |
| Vla | Unknown/<br>Non-study | AK | 144 | U | No | -1.07 (-1.09) | 1132 (1079) | 0.21 | 16.26 | 2.97 | 16 | 18 |
| Vry | Unknown/<br>Non-study | NH | 5 | U | No | -0.95 (-0.94) | 980 (981) | 0.21 | 3.81 | 29.21 | 1 | 11 |
| Vul | BD | NH | 557 | H | Yes | -0.40 (-0.42) | 1240 (1266) | 0.79 | 28.28 | 24.20 | 16 | 30 |
| War | Unknown/<br>Non-study | BD | 3 | U | Yes | -1.45 (-1.46) | 844 (884) | 0.31 | 6.22 | 3.18 | 4 | 8 |
| War | Unknown/<br>Non-study | NH | 397 | U | Yes | 1.23 (1.24) | 844 (884) | 0.00 | 6.22 | 3.18 | 0 | 0 |
| Win | Unknown/<br>Non-study | BD | 21 | U | No | -0.95 (-0.92) | 908 (900) | 0.00 | 4.38 | 1.61 | 0 | 4 |
| Xia | NH | BD | 0 | H | No | -1.20 (-1.19) | 927 (961) | 0.48 | 21.32 | 19.82 | 14 | 18 |
| Xin | NH | BD | 11 | H | No | -1.34 (-1.33) | 1051 (1054) | 0.18 | 11.98 | 22.04 | 8 | 24 |
| Xiu | Unknown/<br>Non-study | BD | 16 | U | No | -1.34 (-1.33) |  | 1.22 | 0.00 | 0.00 | 0 | 0 |
| Yan | KB | AK | 608 | H | Yes | 0.99 (0.98) | 798 (798) | 0.00 | 27.50 | 1.50 | 1 | 1 |
| Yaz | KB | BD | 0 | H | No | -0.92 (-0.92) |  | 1.50 | 1.00 | 19.00 | 0 | 0 |

**Table S2:**
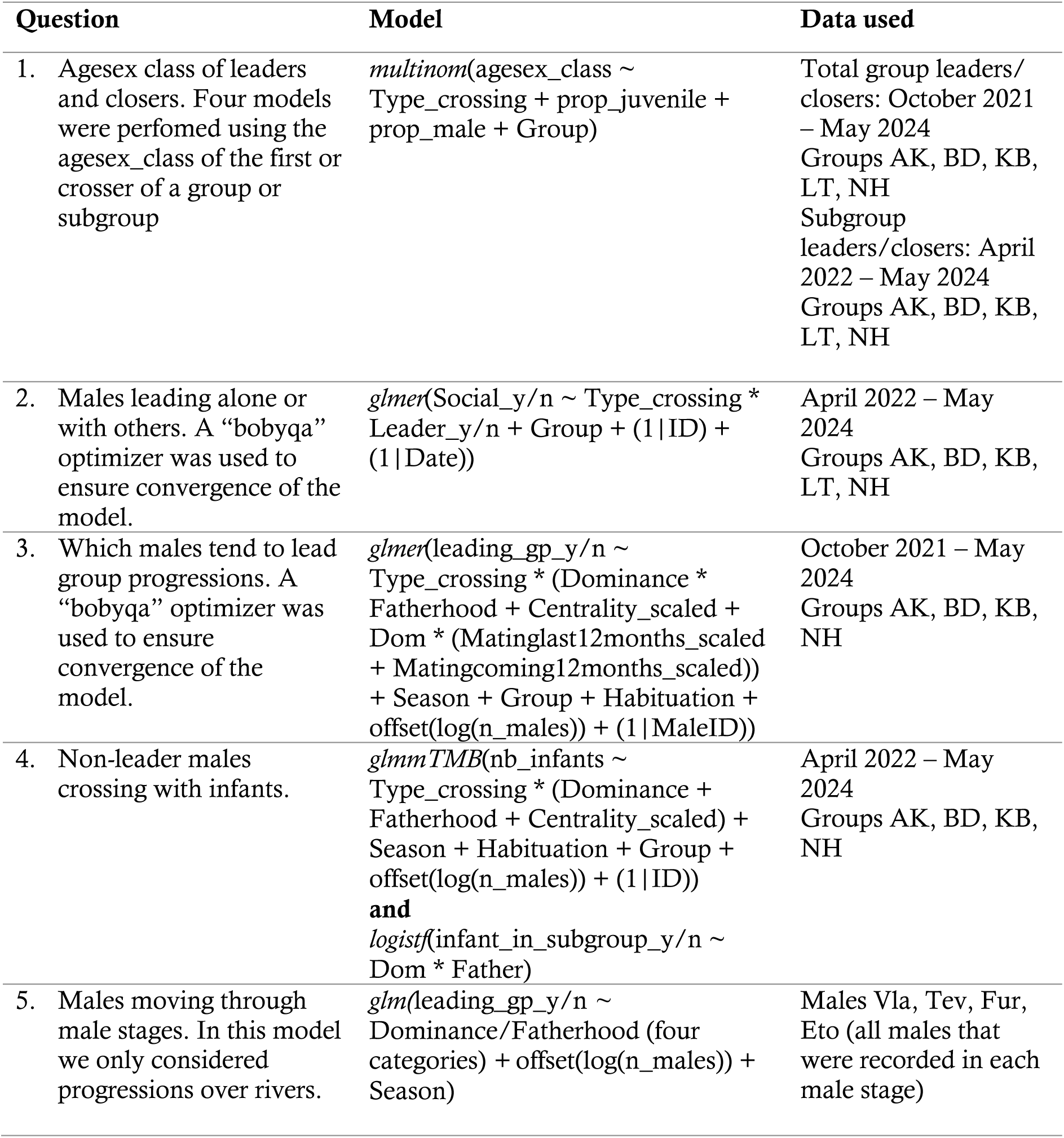
overview of all models performed. Model fit and assumptions were checked using simulation-based residual diagnostics from the DHARMa package (Hartig, 2024).

## Regression tables

In all tables, AM refers to adult males, AF to adult females, and J to juveniles.

**Table 1.**
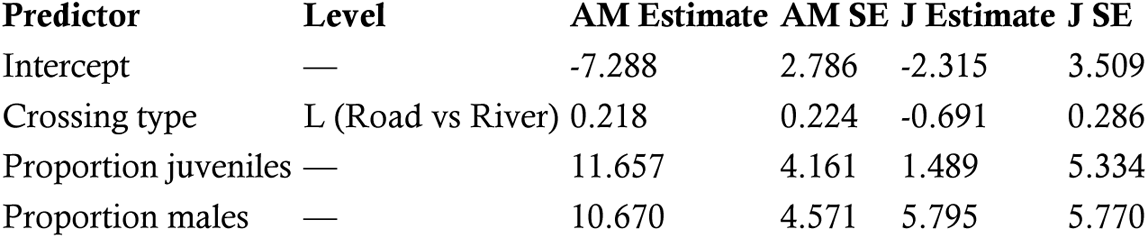
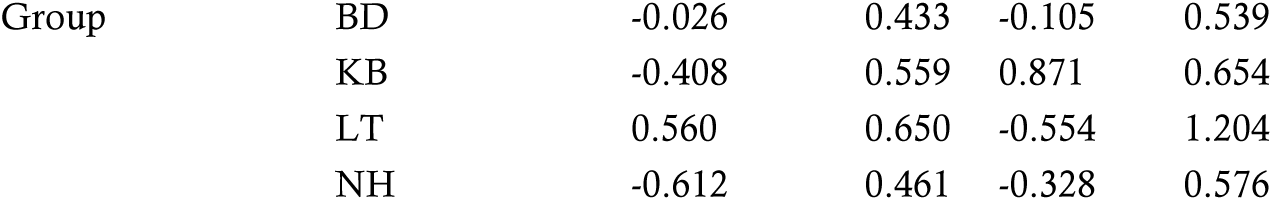
Multinomial logistic regression predicting which age–sex class leads group progressions. Coefficients are log-odds (logits); SE = standard error. Reference category for the outcome is adult females (AF).

| Predictor | Level | AM Estimate | AM SE | J Estimate | J SE |
| --- | --- | --- | --- | --- | --- |
| Intercept | — | -7.288 | 2.786 | -2.315 | 3.509 |
| Crossing type | L (Road vs River) | 0.218 | 0.224 | -0.691 | 0.286 |
| Proportion juveniles | — | 11.657 | 4.161 | 1.489 | 5.334 |
| Proportion males | — | 10.670 | 4.571 | 5.795 | 5.770 |
| Group | BD | -0.026 | 0.433 | -0.105 | 0.539 |
|  | KB | -0.408 | 0.559 | 0.871 | 0.654 |
|  | LT | 0.560 | 0.650 | -0.554 | 1.204 |
|  | NH | -0.612 | 0.461 | -0.328 | 0.576 |

**Table 2.** Pairwise comparisons for leading group progressions. Estimates are log-odds differences from emmeans. Positive values indicate that the first category in the contrast is more likely to lead.

| Contrast | Estimate | SE | df | t | p |
| --- | --- | --- | --- | --- | --- |
| AF – AM | -0.265 | 0.052 | 16 | -5.14 | 0.0003 |
| AF – J | 0.096 | 0.042 | 16 | 2.29 | 0.086 |
| AM – J | 0.361 | 0.048 | 16 | 7.58 | <0.0001 |

**B.** Age–sex differences within each terrain type *Road crossings*
| Contrast | Estimate | SE | df | t | p |
| --- | --- | --- | --- | --- | --- |
| AF – AM | -0.190 | 0.084 | 16 | -2.27 | 0.090 |
| AF – J | 0.018 | 0.074 | 16 | 0.24 | 0.968 |
| AM – J | 0.208 | 0.084 | 16 | 2.49 | 0.059 |

*River crossings*
| Contrast | Estimate | SE | df | t | p |
| --- | --- | --- | --- | --- | --- |
| AF – AM | -0.339 | 0.067 | 16 | -5.04 | 0.0003 |
| AF – J | 0.174 | 0.044 | 16 | 3.98 | 0.0029 |
| AM – J | 0.513 | 0.049 | 16 | 10.45 | <0.0001 |

**C.** Terrain effect within each age–sex class (Road – River)
| Age–sex | Estimate | SE | df | t | p |
| --- | --- | --- | --- | --- | --- |
| AF | -0.002 | 0.0585 | 16 | -0.04 | 0.968 |
| AM | -0.151 | 0.064 | 16 | -2.37 | 0.031 |
| J | 0.154 | 0.050 | 16 | 3.09 | 0.007 |

**Table 3.** Multinomial logistic regression predicting which age–sex class closes group progressions. Coefficients are log-odds; SE = standard error. Reference category for the outcome is adult females (AF).

| Predictor | Level | AM Estimate | AM SE | J Estimate | J SE |
| --- | --- | --- | --- | --- | --- |
| Intercept | — | -8.385 | 4.121 | -2.999 | 3.928 |
| Crossing type | L (Road vs River) | -0.056 | 0.325 | -0.499 | 0.319 |
| Proportion juveniles | — | 9.926 | 5.979 | 3.738 | 5.847 |
| Proportion males | — | 12.526 | 6.478 | -3.057 | 6.343 |
| Group | BD | 1.628 | 0.605 | 1.128 | 0.617 |
|  | KB | 0.395 | 0.755 | 1.174 | 0.756 |
|  | LT | 1.135 | 0.816 | 1.382 | 0.781 |
|  | NH | 0.616 | 0.595 | 1.292 | 0.586 |

Residual deviance = 530.27; AIC = 562.27. Likelihood-ratio tests (Type II)
| Predictor | LR $\chi^2$ | df | p |
| --- | --- | --- | --- |
| Crossing type | 2.94 | 2 | 0.230 |
| Proportion juveniles | 2.86 | 2 | 0.239 |
| Proportion males | 5.48 | 2 | 0.065 |
| Group | 14.28 | 8 | 0.075 |

**Table 4.** Pairwise contrasts for closing group progressions. Estimates are log-odds differences; positive values indicate that the first category in the contrast is more likely to close.

| Contrast | Estimate | SE | df | t | p |
| --- | --- | --- | --- | --- | --- |
| AF – AM | 0.095 | 0.062 | 16 | 1.52 | 0.306 |
| AF – J | 0.116 | 0.065 | 16 | 1.78 | 0.210 |
| AM – J | 0.022 | 0.067 | 16 | 0.32 | 0.945 |

**B.** Age–sex differences within each terrain type. *Road crossings*
| Contrast | Estimate | SE | df | t | p |
| --- | --- | --- | --- | --- | --- |
| AF – AM | 0.074 | 0.106 | 16 | 0.69 | 0.770 |
| AF – J | 0.007 | 0.118 | 16 | 0.06 | 0.998 |
| AM – J | -0.067 | 0.116 | 16 | -0.58 | 0.835 |

*River crossings*
| Contrast | Estimate | SE | df | t | p |
| --- | --- | --- | --- | --- | --- |
| AF – AM | 0.116 | 0.083 | 16 | 1.40 | 0.366 |
| AF – J | 0.225 | 0.070 | 16 | 3.20 | 0.015 |
| AM – J | 0.110 | 0.065 | 16 | 1.67 | 0.245 |

**C.** Terrain effect within each age–sex class (Road – River).
| Age–sex | Estimate | SE | df | t | p |
| --- | --- | --- | --- | --- | --- |
| AF | -0.087 | 0.086 | 16 | -1.01 | 0.327 |
| AM | -0.045 | 0.078 | 16 | -0.57 | 0.576 |
| J | 0.131 | 0.078 | 16 | 1.68 | 0.112 |

| Contrast | Estimate | SE | df | t | p |
| --- | --- | --- | --- | --- | --- |
| AF – AM | 0.397 | 0.138 | 16 | 2.87 | 0.028 |
| AF – J | 0.453 | 0.112 | 16 | 4.03 | 0.003 |
| AM – J | 0.055 | 0.088 | 16 | 0.63 | 0.808 |

**Table 5.**
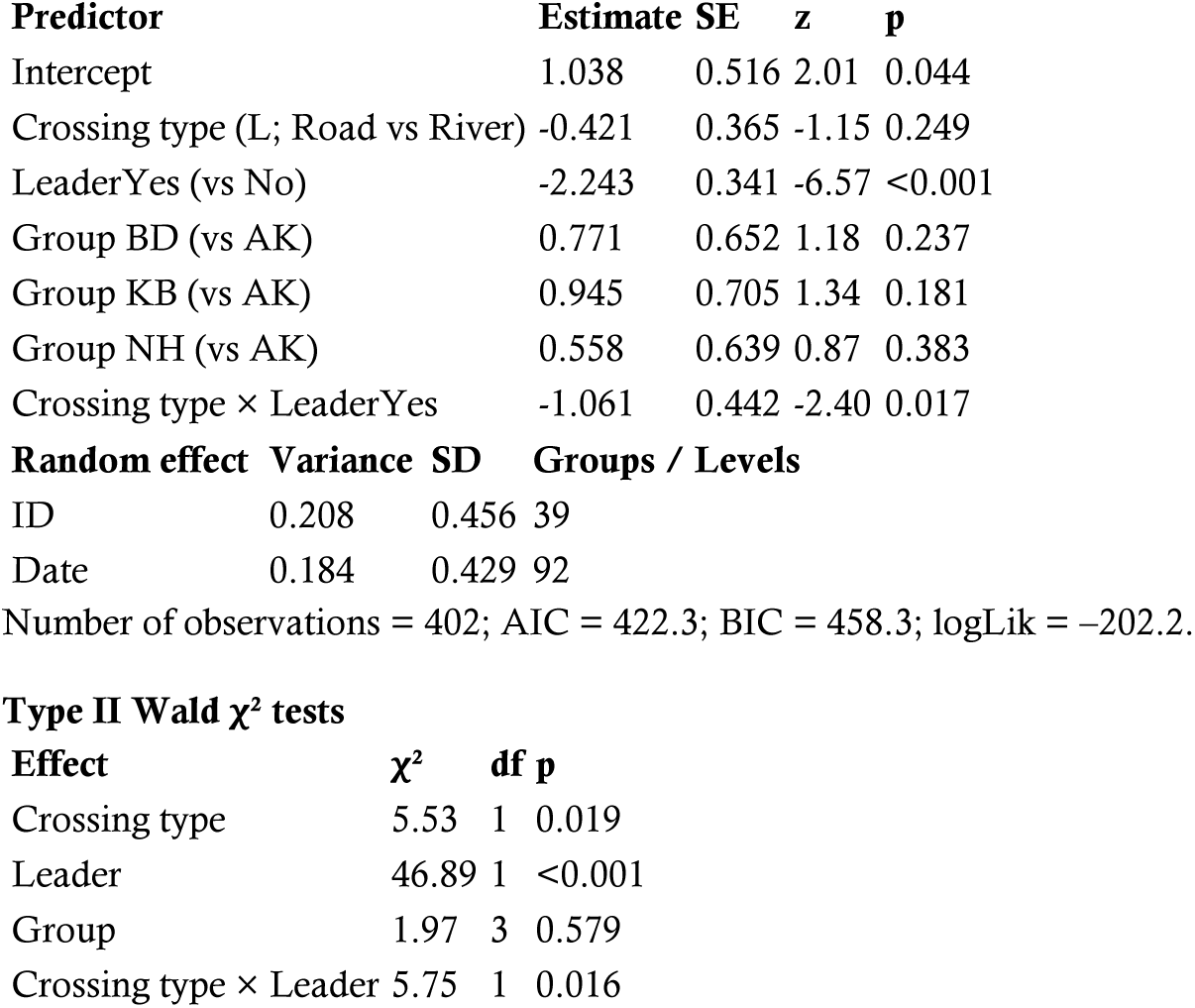
Generalized linear mixed model predicting whether a male crossed with others (within 43 s). Estimates are log-odds; positive values indicate a higher probability of crossing with others.

| Predictor | Estimate | SE | z | p |
| --- | --- | --- | --- | --- |
| Intercept | 1.038 | 0.516 | 2.01 | 0.044 |
| Crossing type (L; Road vs River) | -0.421 | 0.365 | -1.15 | 0.249 |
| LeaderYes (vs No) | -2.243 | 0.341 | -6.57 | <0.001 |
| Group BD (vs AK) | 0.771 | 0.652 | 1.18 | 0.237 |
| Group KB (vs AK) | 0.945 | 0.705 | 1.34 | 0.181 |
| Group NH (vs AK) | 0.558 | 0.639 | 0.87 | 0.383 |
| Crossing type × LeaderYes | -1.061 | 0.442 | -2.40 | 0.017 |
| Random effect | Variance | SD | Groups / Levels |  |
| ID | 0.208 | 0.456 | 39 |  |
| Date | 0.184 | 0.429 | 92 |  |

Type II Wald $\chi^2$ tests
| Effect | $\chi^2$ | df | p |
| --- | --- | --- | --- |
| Crossing type | 5.53 | 1 | 0.019 |
| Leader | 46.89 | 1 | <0.001 |
| Group | 1.97 | 3 | 0.579 |
| Crossing type × Leader | 5.75 | 1 | 0.016 |

**Table 6.** Pairwise contrasts for crossing socially. All estimates are log-odds ratios.

| Contrast | Estimate | SE | z | p |
| --- | --- | --- | --- | --- |
| No – Yes | 2.24 | 0.341 | 6.57 | <0.0001 |

**B.** Leaders vs non-leaders within each terrain.
| Crossing type | Contrast | Estimate | SE | z | p |
| --- | --- | --- | --- | --- | --- |
| Road | No – Yes | 1.49 | 0.486 | 3.08 | 0.0021 |
| River | No – Yes | 2.99 | 0.439 | 6.82 | <0.0001 |

C. Terrain effect within each leadership status (Road – River).
| Leader | Contrast | Estimate | SE | z | p |
| --- | --- | --- | --- | --- | --- |
| No | Road – River | 0.595 | 0.516 | 1.15 | 0.249 |
| Yes | Road – River | 2.096 | 0.626 | 3.35 | 0.0008 |

**Table 7.**
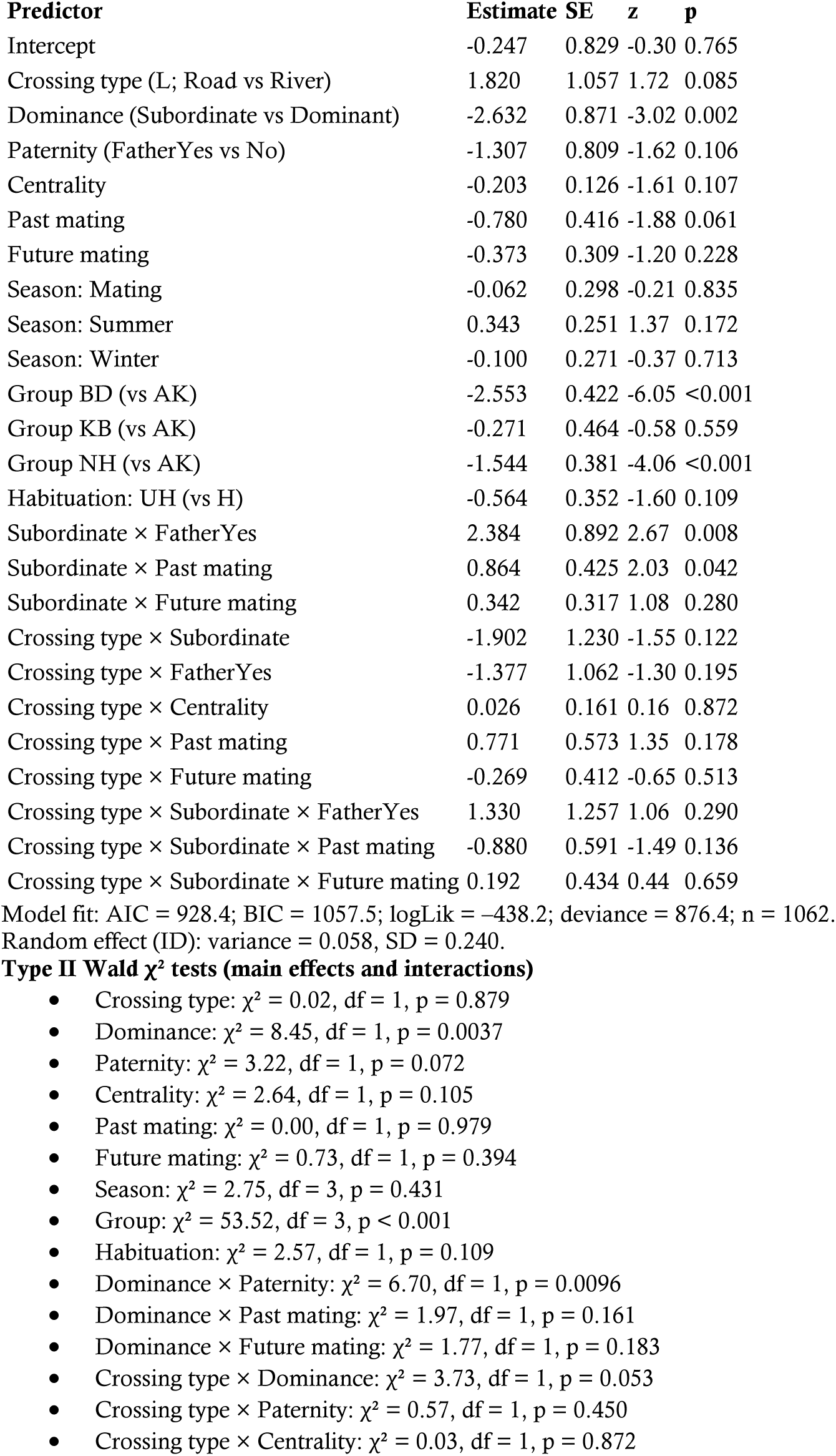
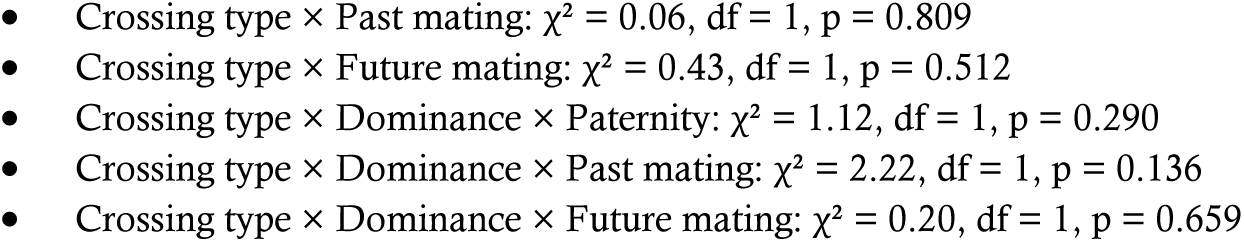
GLMM predicting male probability to lead group progressions. Estimates are log-odds; positive values indicate a higher probability of leading.

**Table 8.**
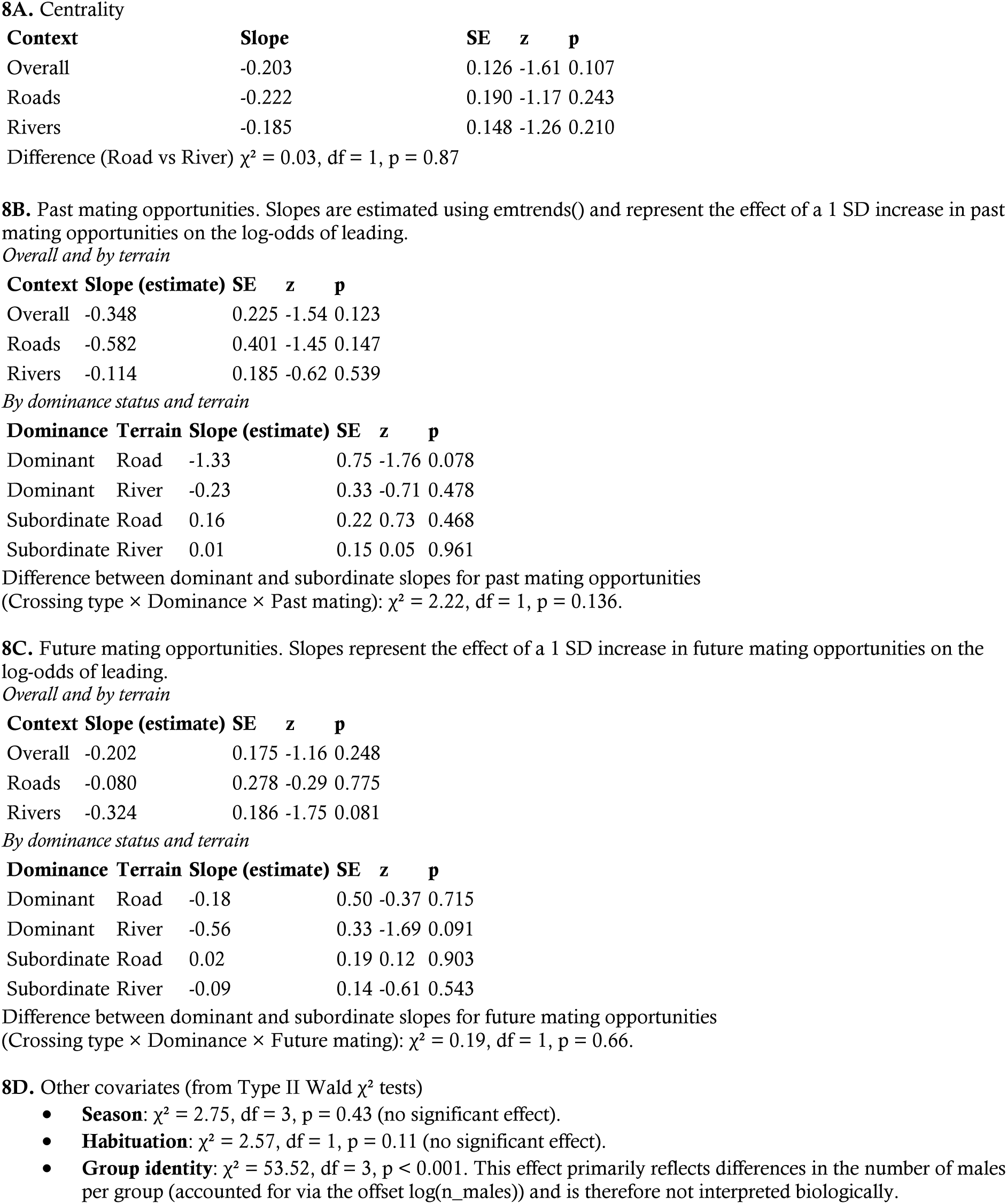
Centrality and mating-opportunity slopes (emtrends). Slopes are from emtrends() and represent the effect of a 1 SD change in the standardized predictor on the log-odds of leading.

## References

Alberts, S. C., Watts, H. E., & Altmann, J. (2003). Queuing and queue-jumping: Long-term patterns of reproductive skew in male savannah baboons, Papio cynocephalus. Animal Behaviour, 65(4), 821–840. 10.1006/anbe.2003.2106

Axelrod, R., & Hamilton, W. D. (1981). The Evolution of Cooperation Published by: American Association for the Advancement of Science The Evolution of Cooperation. Science, 211(4489), 1390–1396.

Baldellou, M., & Henzi, S. P. (1992). Vigilance, predator detection and the presence of supernumerary males in vervet monkey troops. Animal Behaviour, 43(3), 451–461. 10.1016/S0003-3472(05)80104-6

Baldellou, M., Henzi, S. P., & Choe, J.-H. (1992). Vigilance, predator detection and the presence of supernumerary males in vervet monkey troops. Animal Behaviour, 43(3), 123–128. 10.1016/S0003-3472(05)80104-6

Barelli, C., Reichard, U., Boesch, C., & Heistermann, M. (2008). Female white-handed gibbons (*Hylobates lar*) lead group movements and have priority of access to food resources. Behaviour, 145(7), 965–981. 10.1163/156853908784089243

Borgeaud, C., Sosa, S., Bshary, R., Sueur, C., & van De Waal, E. (2016). Intergroup Variation of Social Relationships in Wild Vervet Monkeys: A Dynamic Network Approach. Frontiers in Psychology, 7. 10.3389/fpsyg.2016.00915

Bumann, D., Krause, J., & Rubenstein, D. I. (1997). Mortality risk of spatial positions in animal groups: The danger of being in the front. Behaviour, 134, 1063–1076.

Cheney, D. L., & Seyfarth, R. M. (1990). The representation of social relations by monkeys. Cognition, 37(1-2), 167–196. 10.1016/0010-0277(90)90022-C

Cheney, D., Seyfarth, R., & Smuts, B. (1986). Social Relationships and Social Cognition in Nonhuman Primates Author (*s*): Dorothy Cheney, Robert Seyfarth and Barbara Smuts. Science (New York, N.Y.), 234(4782), 1361–1366.

Clutton-Brock, T. (2009). Cooperation between non-kin in animal societies. Nature, 462(7269), 51–57. 10.1038/nature08366

Farine, D. R., Strandburg-Peshkin, A., Couzin, I. D., Berger-Wolf, T. Y., & Crofoot, M. C. (2017). Individual variation in local interaction rules can explain emergent patterns of spatial organization in wild baboons. Proceedings of the Royal Society B: Biological Sciences, 284(1853), 20162243. 10.1098/rspb.2016.2243

Fichtel, C., Pyritz, L., & Kappeler, P. M. (2011). Coordination of Group Movements in Non-human Primates. 37–56. 10.1007/978-3-642-15355-6

Hall, K. R. L. (1960). Social vigilance behaviour of the chacma baboon, Papio ursinus. Behaviour, 16, 261–294. 10.1163/15685-3960X00188

Hamilton, W. D. (1964). The genetical evolution of social behaviour. II. Journal of Theoretical Biology, 7(1), 17–52. 10.1016/0022-5193(64)90039-6

Hamilton, W. D. (1971). Geometry for the selfish herd. Journal of Theoretical Biology, 31(2), 295–311. 10.1016/0022-5193(71)90189-5

Henzi, S. P., & Lucas, J. W. (1980). Observations on the Inter-Troop Movement of Adult Vervet Monkeys (*Cercopithecus aethiops*). Folia primatol., 33, 220–235.

Hockings, K. J. (2011). Behavioral Flexibility and Division of Roles in Chimpanzee Road-Crossing. In T. Matsuzawa, T. Humle, & Y. Sugiyama (Red.), The Chimpanzees of Bossou and Nimba (pp. 221–229). Springer Japan. 10.1007/978-4-431-53921-6_24

Hockings, K. J., Anderson, J. R., & Matsuzawa, T. (2006). Road crossing in chimpanzees: A risky business. Current Biology, 16(17), 668–670. 10.1016/j.cub.2006.08.019

Kavanagh, M. (1980). Invasion of the Forest By an African Savannah Monkey: Behavioural Adaptations. Behaviour, 73(3-4), 238–260. 10.1163/156853980X00258

King, A. J., Douglas, C. M. S., Huchard, E., Isaac, N. J. B., & Cowlishaw, G. (2008). Dominance and Affiliation Mediate Despotism in a Social Primate. Current Biology, 18(23), 1833–1838. 10.1016/j.cub.2008.10.048

Lukas, D., & Clutton-Brock, T. (2018). Social complexity and kinship in animal societies. Ecology Letters, 21(8), 1129–1134. 10.1111/ele.13079

Mercier, S., Neumann, C., van De Waal, E., Chollet, E., Meric De Bellefon, J., & Zuberbühler, K. (2017). Vervet monkeys greet adult males during high-risk situations. Animal Behaviour, 132, 229–245. 10.1016/j.anbehav.2017.07.021

Minkner, M. M. I., Young, C., Amici, F., McFarland, R., Barrett, L., Grobler, J. P., Henzi, S. P., & Widdig, A. (2018). Assessment of Male Reproductive Skew via Highly Polymorphic STR Markers in Wild Vervet Monkeys, *Chlorocebus pygerythrus*. Journal of Heredity. 10.1093/jhered/esy048

Muroyama, Y. (1998). Reciprocal Altruistic Behaviour in Non-Human Primates. Primate Research, 14(3), 165–178. 10.2354/psj.14.165

Neumann, C. (2016). socialindices—A (not entirely) brief tutorial to quantify social bonds with R [Software]. Neumann, C., & Kulik, L. (2024). EloRating—A brief tutorial (Versie 0.46.18) [Software].

Raihani, N. J., & Bshary, R. (2011). Resolving the iterated prisoner’s dilemma: Theory and reality: Resolving the IPD. Journal of Evolutionary Biology, 24(8), 1628–1639. 10.1111/j.1420-9101.2011.02307.x

Rhine, R. J., Forthman, D. L., Stillwell-Barnes, R., Westlund, B. J., & Westlund, H. D. (1981). Movement patterns of yellow baboons (*Papio cynocephalus*): Sex differences in juvenile development toward adult patterns. American Journal of Physical Anthropology, 55(4), 473–484. 10.1002/ajpa.1330550408

Rhine, R. J., & Westlund, B. J. (1981). Adult Male Positioning in Baboon Progressions: Order and Chaos Revisited. Folia Primatologica, 35, 77–116.

Ripley, B., & Venables, W. (2016). Package ‘nnet’. R package version, 7(3-12), 700.

Roberts, G. (1998). Competitive altruism: From reciprocity to the handicap principle. In Proceedings of the Royal Society B: Biological Sciences (Vol. 265, Nummer 1394, pp. 427–431). 10.1098/rspb.1998.0312

Romenskyy, M., Herbert-Read, J. E., Ioannou, C. C., Szorkovszky, A., Ward, A. J. W., & Sumpter, D. J. T. (2020). Quantifying the structure and dynamics of fish shoals under predation threat in three dimensions. Behavioral Ecology, 31(2), 311–321. 10.1093/beheco/arz197

Rowell, T. E. (1966). Forest living baboons in Uganda. Journal of Zoology, 149(3), 344–364. 10.1111/j.1469-7998.1966.tb04054.x

Seyfarth, R. M., & Cheney, D. L. (2000). Social Awareness in Monkeys. American Zoology, 40, 902–909.

Stephan, C., & Zuberbühler, K. (2016). Persistent Females and Compliant Males Coordinate Alarm Calling in Diana Monkeys. Current Biology, 26(21), 2907–2912. 10.1016/j.cub.2016.08.033

Stevens, J. R., Cushman, F. A., & Hauser, M. D. (2005). Evolving the Psychological Mechanisms for Cooperation. Annual Review of Ecology, Evolution, and Systematics, 36(1), 499–518. 10.1146/annurev.ecolsys.36.113004.083814

Stojan-Dolar, M., & Heymann, E. W. (2010). Vigilance in a cooperatively breeding primate. International Journal of Primatology, 31(1), 95–116. 10.1007/s10764-009-9385-7

Stoltz, P., & Saayman, G. S. (1970). Ecology and behaviour of baboons in the Northern Transvaal. Annals of the Transvaal Museum, 26(99-143).

Sueur, C., & Petit, O. (2008). Organization of Group Members at Departure Is Driven by Social Structure in Macaca. International Journal of Primatology, 29(4), 1085–1098. 10.1007/s10764-008-9262-9

Suire, A., Kunita, I., Harel, R., Crofoot, M., Mutinda, M., Kamau, M., Hassel, J. M., Murray, S., Kawamura, S., & Matsumoto-Oda, A. (2023). Estimating individual exposure to predation risk in group-living baboons, *Papio anubis*. PLOS ONE, 18(11), e0287357. 10.1371/journal.pone.0287357

Teichroeb, J. A., White, M. M. J., & Chapman, C. A. (2015). Vervet (*Chlorocebus pygerythrus*) Intragroup Spatial Positioning: Dominants Trade-Off Predation Risk for Increased Food Acquisition. International Journal of Primatology, 36(1), 154–176. 10.1007/s10764-015-9818-4

Tellier, M., Druelle, F., Cibot, M., Baruzaliire, J., Sabiiti, T., & McLennan, M. R. (2025). Running the Risk: Road-Crossing Behavior in Wild Chimpanzees (*Pan troglodytes*) in an Anthropogenic Habitat in Uganda. American Journal of Primatology, 87(2), e70000. 10.1002/ajp.70000

van Noordwijk, M. A., & van Schaik, C. P. (1988). Male Careers in Sumatran Long-Tailed Macaques (*Macaca Fascicularis*). Behaviour, 24–43.

van Schaik, C. P., Bshary, R., Wagner, G., & Cunha, F. (2021). Male anti-predator services in primates as costly signalling? A comparative analysis and review. in review.

van Schaik, C. P., Bshary, R., Wagner, G., & Cunha, F. (2022). Male anti-predation services in primates as costly signalling? A comparative analysis and review. Ethology, 128(1), 1–14. 10.1111/eth.13233

van Schaik, C. P., & Paul, A. (1996). Male care in primates: Does it ever reflect paternity? Evolutionary Anthropology, 5(5), 152–156. 10.1002/(SICI)1520-6505(1996)5:5<152::AID-EVAN3>3.0.CO;2-H

van Schaik, C. P., & van Noordwijk, M. A. (1989). The special role of male Cebus monkeys in predation avoidance and its effect on group composition. Behavioral Ecology and Sociobiology, 24(5), 265–276. 10.1007/BF00290902

Watts, D. P. (1994). The Influence of Male Mating Tactics on Habitat Use in Mountain Gorillas (*Gorilla gorilla beringel*). Primates, 35(1), 35–47.

Weingrill, T., Willems, E. P., Krützen, M., & Noë, R. (2011). Determinants of Paternity Success in a Group of Captive Vervet Monkeys (*Chlorocebus aethiops sabaeus*). International Journal of Primatology, 32, 415–429. 10.1007/s10764-010-9478-3

West, S. A., Griffin, A. S., & Gardner, A. (2007). Evolutionary Explanations for Cooperation. Current Biology, 17(16), 661–672. 10.1016/j.cub.2007.06.004

Young, C., McFarland, R., Barrett, L., & Henzi, S. P. (2017). Formidable females and the power trajectories of socially integrated male vervet monkeys. Animal Behaviour, 125, 61–67. 10.1016/j.anbehav.2017.01.006

Zahavi, A. (1975). Mate selection—A selection for a handicap. Journal of Theoretical Biology, 53(1), 205–214. 10.1016/0022-5193(75)90111-3

